# Exogenous BMP9 therapy ameliorates primary graft dysfunction post lung transplantation

**DOI:** 10.1101/2025.09.30.679530

**Authors:** Blake Gill, Zhenxiao Tu, Logan Langerude, Amir Emtiazjoo, Ashish K Sharma, Carl Atkinson

## Abstract

Primary graft dysfunction (PGD) is a leading cause of early mortality after lung transplantation and is driven by ischemia-reperfusion injury (IRI), which destabilizes the pulmonary endothelium. Bone morphogenetic protein 9 (BMP9), a vascular quiescence factor of the TGF superfamily, maintains endothelial homeostasis through ALK-1/BMPR2-mediated signaling. Herein, this work identified a previously unrecognized suppression of BMP9 signaling during early reperfusion in a murine orthotopic lung transplant model. Allograft transcriptomic profiling revealed rapid downregulation of BMP9 pathway components, including Bmpr2, Acvrl1, Smad5, and Eng. Recombinant BMP9 administered at reperfusion enhanced Id1 expression in pulmonary endothelium, reduced neutrophil infiltration, preserved vascular barrier function, and improved oxygenation. In vitro, BMP9 attenuated proinflammatory cytokine release following simulated cold ischemia-reperfusion injury in human lung endothelial cells. Plasma from lung transplant recipients, regardless of PGD status, had impaired BMP9-induced ID1 expression in endothelial cells, with the most profound suppression observed in patients with interstitial lung disease. These findings implicate transient loss of BMP9 signaling as a key feature of transplant-induced vascular injury and support the therapeutic potential of BMP9 supplementation to restore endothelial homeostasis and improve early graft outcomes.

## INTRODUCTION

Lung transplantation (LTx) remains the only definitive treatment for patients with end-stage pulmonary disease, yet early graft failure due to primary graft dysfunction (PGD) significantly limits its success (1). PGD presents as hypoxemia and non-cardiogenic pulmonary edema, reflecting diffuse alveolar injury, within the first 72hr post-implantation (2). Ischemia-reperfusion injury (IRI) is the primary driver of PGD where the abrupt alterations in temperature and mechanical stressors provoke pulmonary endothelial activation, innate immune recruitment, and alveolar-capillary barrier dysfunction (3–5). No targeted therapies exist to prevent or treat PGD, the leading cause of morbidity and mortality, underscoring the need to better define molecular mechanisms underlying early allograft injury (6).

The pulmonary endothelium plays a central role in PGD pathogenesis (4). Under physiological conditions, lung microvascular endothelial cells maintain barrier integrity, regulate coagulation, and suppress leukocyte adhesion through a quiescent, anti-inflammatory phenotype. IRI triggers endothelial cells to upregulate adhesion molecules, lose junctional stability, degrade the glycocalyx, and secrete pro-inflammatory mediators resulting in increased immunogenicity, vascular permeability, and alveolar edema (7–10). While endothelial dysfunction is recognized as a driver of PGD, the upstream regulatory pathways governing this transition remain poorly understood (11, 12).

Bone morphogenetic protein 9 (BMP9), encoded by growth differentiation factor 2 (Gdf2), is predominantly produced by hepatocytes before its secretion into the circulatory system (13). As with other transforming growth factor (TGF) superfamily members, BMP9 binds a heterotetramer of type I and type II receptors. In the pulmonary microvasculature, activin receptor-like kinase 1 (ALK-1) and bone morphogenetic protein receptor 2 (BMPR2) are the primary type I and II receptors, respectively (14–16). Upon ligand binding, BMP9 signals through ALK-1/BMPR2 to induce phosphorylation of small mothers against decapentaplegic proteins (SMAD) 1/5/9, which then interact with SMAD4 to translocate to the nucleus to regulate target gene transcription (13, 17, 18). This signaling axis promotes endothelial quiescence, enhances intercellular junction stability, and suppresses pro-inflammatory gene expression, thereby maintaining microvascular barrier function (19–22). Loss-of-function receptor mutations and/or deficiencies in BMP9 signaling underlying diseases such as pulmonary arterial hypertension (PAH) and hereditary hemorrhagic telangiectasia (HHT) 1 and 2, which are characterized by endothelial instability, arteriovenous malformations, excessive angiogenesis, and vascular leak (23–27).

Beyond chronic vascular diseases, recent clinical studies have identified reductions in circulating BMP9 levels in critically ill patients with sepsis and acute respiratory distress syndrome (ARDS), conditions similarly characterized by pulmonary endothelial dysfunction and capillary leak (22, 28, 29). These observations raise the possibility that BMP9 deficiency, or signaling impairment, represents a broader mechanism of endothelial barrier failure in acute lung injury syndromes. Despite these insights, the role of BMP9 signaling in the early stages of LTx-associated PGD remains undefined. Whether IRI disrupts BMP9 signaling in the pulmonary endothelium, and contributes to early microvascular dysfunction is unknown. To address this, targeted transcriptomic profiling of murine orthotopic LTx harvested at early (6-hour) and later (24-hour) post-reperfusion intervals were performed. Using NanoString nCounter analysis, we assessed temporal changes in endothelial associated signaling pathways, with a specific focus on BMP signaling components. Using in vitro and in vivo models we show that BMP9-BMPR2 signaling is downregulated early post transplantation and that restoring signaling improves outcomes. Taken together, our study identifies a tractable immuno-vascular pathway with direct therapeutic implications for LTx.

## Results

### BMP9 signaling is disrupted early after lung transplantation

A NanoString nCounter analysis was performed on RNA isolated from mice that had received a fully allogeneic left orthotopic LTx (Figure 1A). We compared nCounter gene expression analysis on allografts harvested at either 6 or 24 hrs post-LTx to normal Balb/c donor lungs. Bi-Plot analysis of the normalized count PCA-X model data revealed that murine normal, 6 hr and 24 hr post-LTx lungs have distinctively different transcript signatures (Figure 1B). Hierarchal clustering analysis (HCA), represented by a dendrogram calculated with Ward’s, corroborated this result (Supplemental Figure 1A). Normal vs 6 hr post-LTx lungs data can be found in Supplemental Figure 1B. Differential expression analysis identified *bmpr2* as one of the most significantly downregulated genes (Figure 1C & 1D). Notably, proinflammatory mediators were upregulated following LTx. We further analyzed other BMP signaling genes present in the nCounter panel and found Acvrl1, Eng, Bmpr1a, and Smad5, important factors in the BMP signaling cascade that were similarly significantly reduced (Figure 1E-1H). To validate these expression changes, RT-qPCR was employed and confirmed that transcript expression of *acvrl1* and *bmpr2* was significantly reduced in lung tissues at 24 hr post-LTx (Supplementary Figure 2A & 2B). Interestingly, the well described BMP9 microvascular target gene, ID1, was not significantly different in the NanoString analysis of bulk lung tissue (Figure 1I), but it was reduced at 24 hr, as shown by RT-qPCR (Supplementary Figure 2C). Other genes involved in the BMP signaling axis, Acvr1, Acvr2a/b, Smad4, and Id2, detected in the NanoString nCounter experiment, were also shown to be differentially regulated post-LTx (Supplemental Figure 3A-3E).

**Figure 1.**
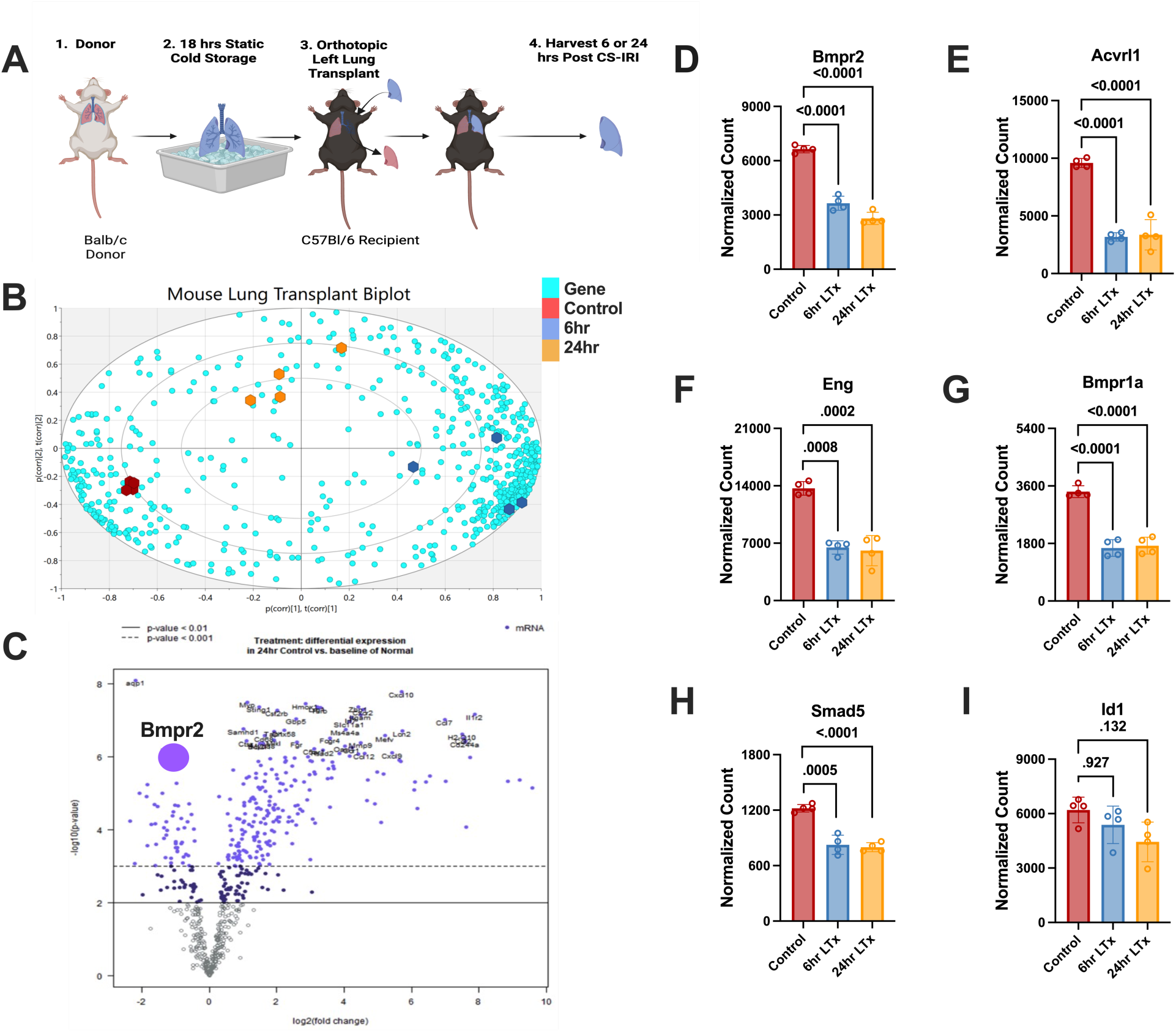
Ischemia-reperfusion injury rapidly suppresses pulmonary BMP9 signaling following lung transplantation. (**A**) Schematic of the fully allogeneic murine left lung transplantation model. Donor lungs from *Balb/c* mice were subjected to 18 hours of cold storage, followed by orthotopic transplantation into *C57BL/6* recipients. Grafts were harvested at 6 and 24 hours post-reperfusion for analysis. (**B**) SIMCA software was used in conjunction with the nCounter NanoString normalized mRNA count data to create a Bi-Plot from the PCA-X model to visualize the relationship between gene expression (loadings) and experimental groups/replicates (scores). (**C**) nSolver generated volcano plot depicting differential gene expression between normal donor lungs and 24-hour post-transplant grafts, highlighting a marked downregulation of Bmpr2, a central receptor in the BMP9 signaling pathway. (**D–G**) Expression of canonical BMP9 receptor components, including Bmpr2 (**D**), Acvrl1 (ALK-1) (**E**), Eng (**F**), and Bmpr1a (**G**), was significantly reduced as early as 6 hours post-transplant and remained suppressed at 24 hours (n=4/group). (**H**) Expression of Smad5, a downstream BMP9 effector, was similarly decreased following transplantation. (**I**) Transcript levels of the BMP9 target gene Id1 were not significantly different at 6 or 24 hours in bulk lung tissue, suggesting cell-specific effects or dilution from non-endothelial compartments. Data represent mean ± SEM; statistical comparisons were performed using one-way ANOVA with Tukey’s post hoc correction. P-values are shown where significant.

### Exogenous BMP9 restores BMP signaling and reduces pro-inflammatory cytokine release

Given our in vivo findings, we next sought to dissect BMP9 signaling specifically within the lung microvascular endothelium using an established in vitro simulated cold storage ischemia-reperfusion injury (CS-IRI) model. Primary human lung microvascular endothelial cells (HMVEC-L) were subjected to CS-IRI, and expression of key BMP9 pathway components (ALK1, BMPR2, ID1) was assessed (Figure 2A-2D). RT-qPCR revealed that CS-IRI significantly reduced BMPR2 and ID1 transcript levels at both 18- and 24-hours post-CS-IRI (Figure 2C & 2D).

**Figure 2.**
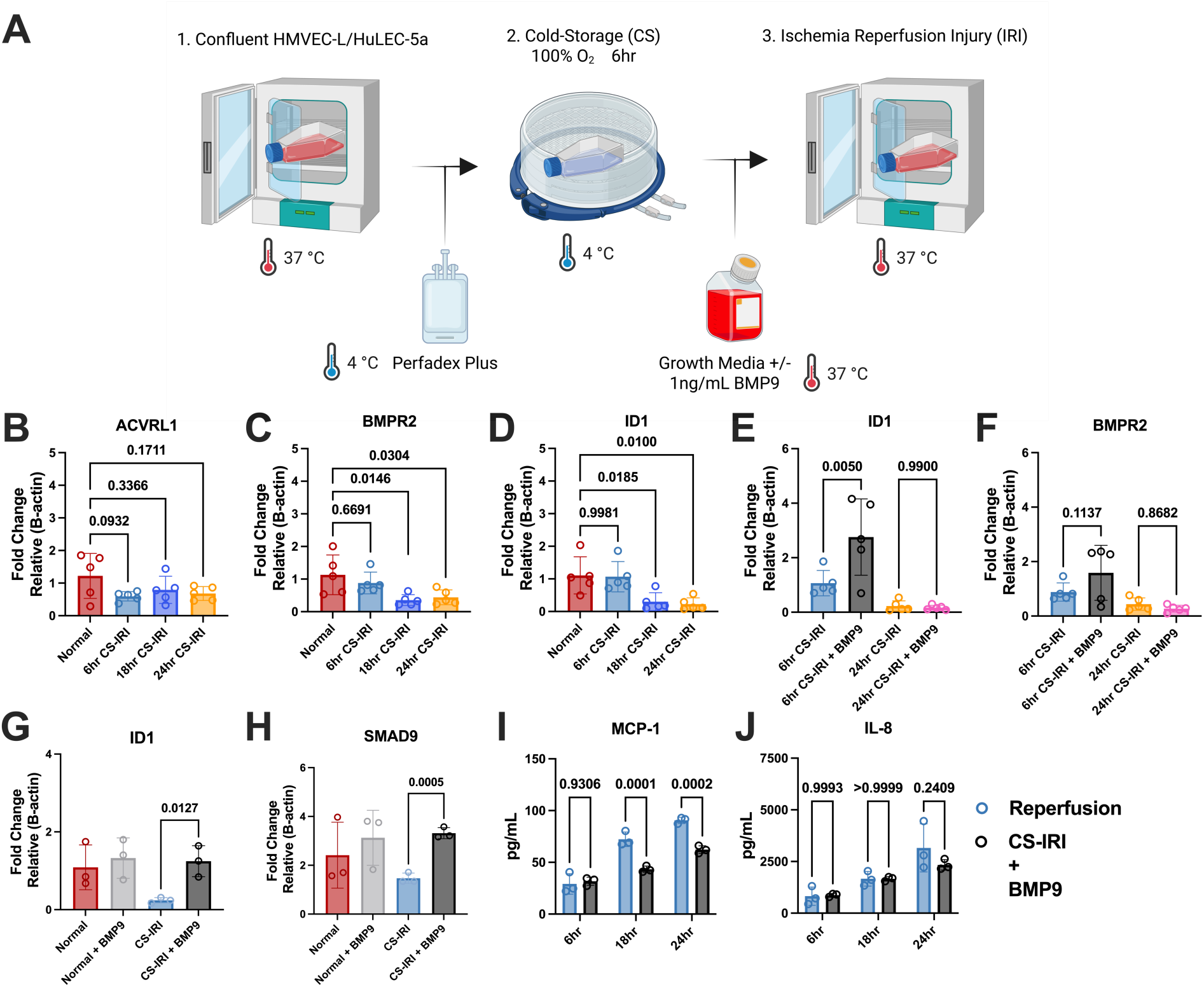
Exogenous BMP9 preserves endothelial BMP signaling and reduces inflammatory responses following cold-storage and ischemia-reperfusion injury (IRI) in vitro. **(A)** Schematic of the in vitro cold-storage (CS) and IRI model using confluent HMVEC-L or HuLEC-5a cells. Cells underwent 6 hours of cold storage in Perfadex Plus at 4°C, followed by reperfusion under normoxic conditions with or without BMP9 (1 ng/mL) supplementation. Samples were harvested at 6, 18, and 24 hours post-reperfusion. **(B–C)** RT-qPCR analysis of *acvrl1* and *bmpr2* expression showed significant reductions following CS/IRI, partially restored by exogenous BMP9. **(D–E)** BMP9 supplementation rescued ID1 expression in both time- and dose-dependent manners, indicating preserved downstream BMP signaling. **(F–G)** BMP9 treatment increased *id1* and *smad9* expression following IRI, consistent with enhanced canonical BMP pathway activation. **(H–I)** Cytokine Elisa assays demonstrated that BMP9 supplementation significantly reduced MCP-1 secretion at 18 and 24 hours post-reperfusion, whereas IL-8 levels were unaffected. Data shown are represented by mean ± SEM. Parts B, C, D, E, F, H, and I statistical comparisons were performed using one-way ANOVA with Tukey’s post hoc correction. Unpaired t-tests were performed for parts F and G. P-values as shown.

To determine whether BMP9 supplementation could restore signaling, exogenous BMP9 was administered at the onset of reperfusion. BMP9 treatment significantly upregulated ID1 expression as early as 6 hours post-reperfusion compared to vehicle-treated controls (Figure 2E), although receptor expression (BMPR2) was not affected (Figure 2F). These findings were validated in an independent pulmonary endothelial cell line, HuLEC-5a, which also showed significant BMP9-mediated upregulation of ID1 (Figure 2G) and SMAD9 (Figure 2H).

Given the protective role of BMP9 in preserving endothelial quiescence, we next assessed whether BMP9 treatment could attenuate endothelial inflammatory activation post-CS-IRI. BMP9-treated HuLEC-5a cells demonstrated significantly reduced secretion of monocyte chemoattractant protein 1 (MCP-1) at both 18- and 24-hours post-reperfusion (Figure 2I). No differences were observed in interleukin-8 (IL-8) secretion (Figure 2J).

Collectively, these findings demonstrate that BMP9 supplementation restores downstream BMP9 signaling and reduces endothelial proinflammatory responses following simulated reperfusion injury.

### Exogenous BMP9 ameliorates lung transplant ischemia-reperfusion injury

To assess the therapeutic potential of BMP9 in mitigating lung transplant-associated injury, we utilized a murine fully allogeneic orthotopic left lung transplant model incorporating prolonged cold storage to induce lung transplant ischemia reperfusion injury (IRI) (Figure 3A). Consistent with our in vitro experiments, recombinant human BMP9 was administered immediately upon reperfusion. BMP9 supplementation significantly improved lung function, as measured by PaO₂, at both 6 and 24 hours post-transplantation compared to vehicle-treated controls (Figure 3B). Histological assessment revealed reduced lung injury in BMP9-treated mice at both timepoints, while administration of a BMP9-blocking antibody exacerbated histological injury (Figure 3C). To determine whether BMP9 modulates early immune infiltration post-transplant, we quantified allograft neutrophil infiltration via myeloperoxidase (MPO) staining at 6 and 24 hours post-LTx. Representative immunofluorescence images illustrate MPO⁺ cells across treatment groups (Figures 4A). BMP9 treatment significantly reduced MPO⁺ neutrophils in the allograft at 6 hours, while anti-BMP9 treatment exacerbated neutrophil infiltration at 24 hours post-transplant (Figure 4B). To assess endothelial injury, serum levels of soluble von Willebrand Factor (svWF), a biomarker of endothelial activation and damage, were measured by ELISA. BMP9-treated mice exhibited significantly lower circulating svWF levels at 24 hours compared to controls (Figure 4C), consistent with reduced endothelial injury. Together, these findings suggest that BMP9 treatment attenuates early neutrophil infiltration and preserves endothelial integrity following lung transplantation.

**Figure 3.**
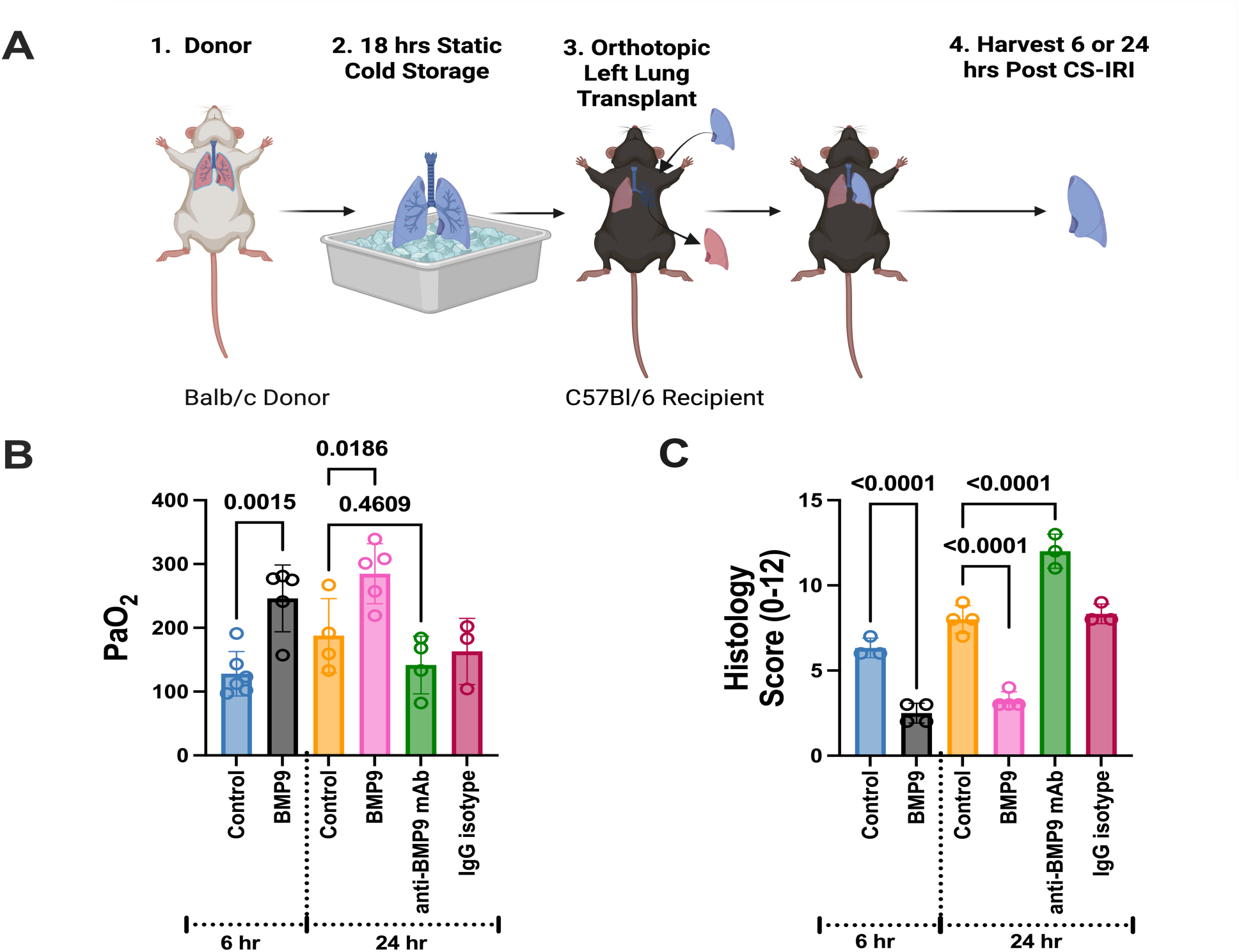
Exogenous BMP9 protects against primary graft dysfunction following lung transplantation. (**A**) Experimental workflow: Balb/c donor lungs were harvested, stored for 18 hours at 4 °C, and transplanted into C57BL/6 recipients. Post-transplant recipients received BMP9, anti-BMP9 blocking antibody, or vehicle control, and grafts were harvested at 6 and 24 hours post cold storage ischemia-reperfusion injury (CS-IRI). (**B**) PaO_2_ revealed that BMP9 treatment significantly improved lung function at both timepoints. (**C**) Lung histology scores (0–12 composite injury scale) revealed marked reduction in injury with BMP9 supplementation at both 6 and 24 hours, whereas anti-BMP9 blockade significantly worsened graft injury compared with vehicle controls. Data are presented as mean ± SEM with individual replicates; statistical significance determined by one-way ANOVA with multiple comparisons (P-values as shown).

**Figure 4.**
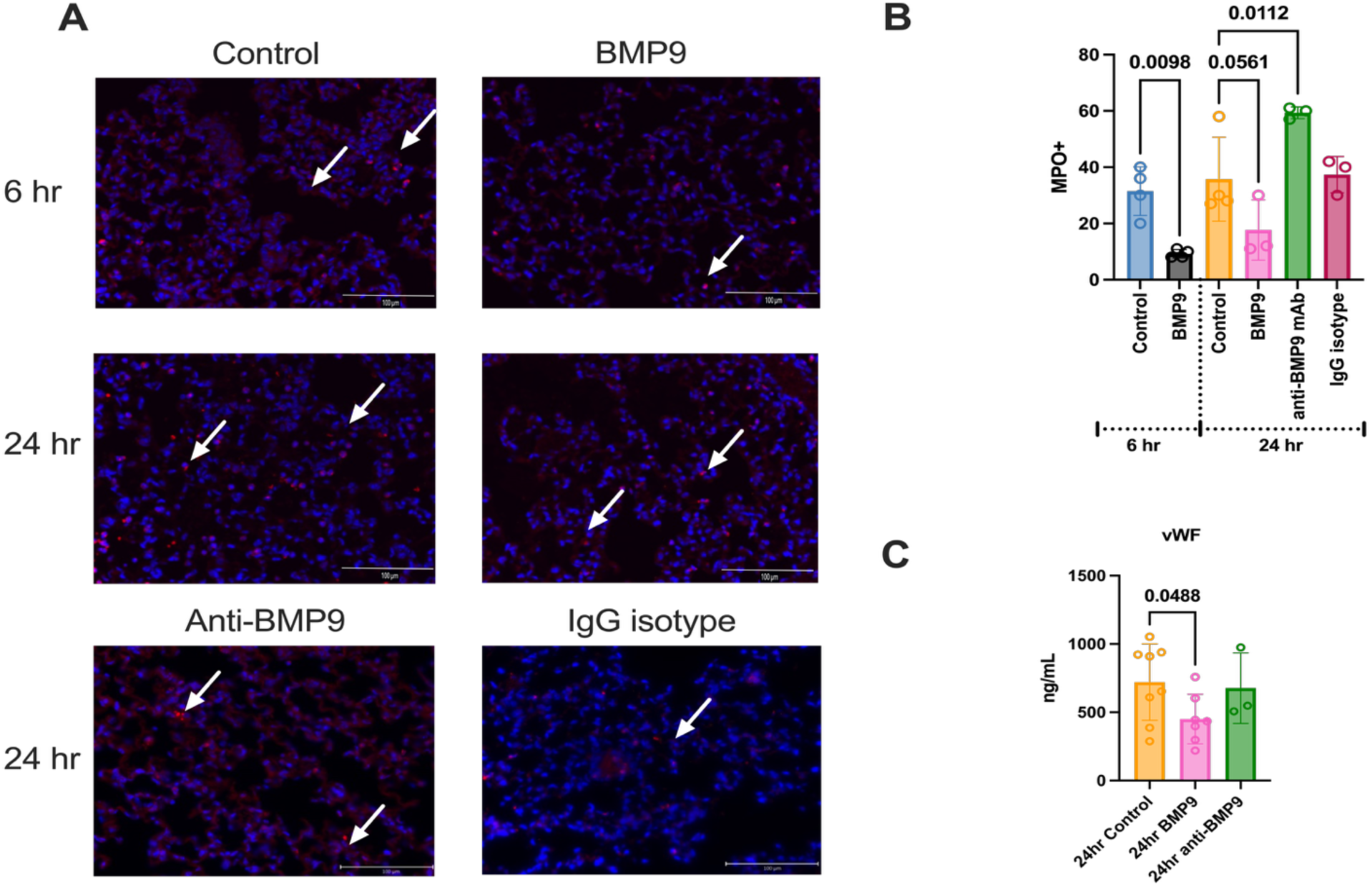
BMP9 attenuates neutrophil infiltration and endothelial injury following murine lung transplantation. (**A**) Representative immunofluorescence staining for myeloperoxidase (MPO, red) in lung sections harvested 24 hours post–cold storage ischemia-reperfusion injury (CS IRI). Nuclei are counterstained with DAPI (blue). BMP9-treated lungs exhibited reduced MPO+ neutrophil infiltration compared with control and anti-BMP9–treated groups. Scale bars = 100 μm. (**B**) Quantification of MPO+ cells per five high-power fields (HPF) demonstrated significantly fewer neutrophils in BMP9-treated recipients at 6hours, whereas anti-BMP9 blockade resulted in increased infiltration at 24 hours. (**C**) Serum von Willebrand Factor (vWF) concentrations at 24 hours post-CS-IRI, reflecting endothelial injury, were significantly reduced in BMP9-treated animals. Data in **B** are shown as mean ± SEM with individual replicates; P-values from one-way ANOVA with multiple comparisons are indicated. Data in **C** are shown as mean ± SEM with individual replicates and an unpaired t-test was performed with the P-value indicated.

### BMP9 therapy acutely enhances vascular Id1 expression

In the lung, BMP9 primarily acts on vascular endothelial cells. Given this specificity, RNAscope for Id1 and Bmpr2 was performed. Representative RNAscope images of Id1 and Bmpr2 within normal, end of cold storage (CS) and the transplanted lung amongst the experimental groups and timepoints are shown (Figure 5A & 5B). At 6 hours post-transplant, BMP9-treated lungs exhibited significantly increased vascular Id1 expression compared to controls (Figure 5A & 5C), with no difference observed at 24 hours. RNAscope analysis of Bmpr2 revealed stable transcript levels across groups and timepoints (Figure 5B & 5D), indicating that enhanced Id1 expression is driven by increased ligand availability rather than receptor up regulation. Interestingly, in vitro BMP9 stimulation increased BMPR2 protein despite stable mRNA, suggesting post-transcriptional regulation (Figures 5E & 5F). These findings demonstrate that BMP9 therapy selectively activates endothelial signaling acutely post-transplant, reinforcing vascular homeostasis through *id1* induction.

**Figure 5.**
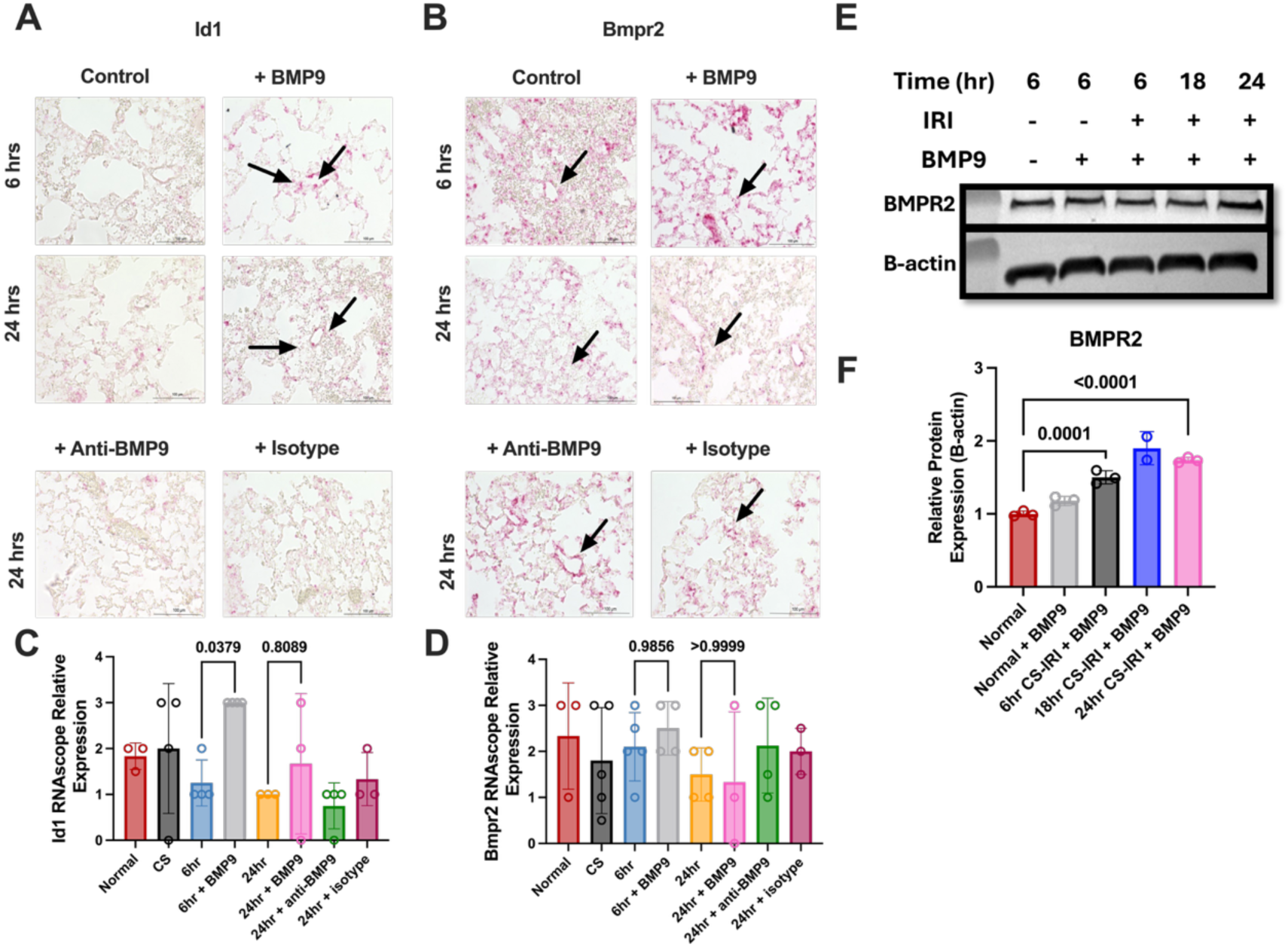
BMP9 regulates downstream BMP signaling targets in transplanted lungs. (**A–B**) Representative RNAscope images demonstrating pulmonary expression of Id1 (**A**) and Bmpr2 (**B**) across experimental groups: Normal, Cold Storage (CS), 6-hours post-transplant, 6-hour post-transplant + BMP9, 24-hour post-transplant, 24-hour post-transplant + BMP9, 24-hour post-transplant + anti-BMP9, and 24-hour post-transplant + isotype control. Positive staining (pink, arrows) highlights transcript localization in alveolar regions. Scale bars = 100 μm. (**C–D**) Quantification of RNAscope relative expression confirmed significant reduction of Id1 at 6 hours post-transplant, with restoration by BMP9 supplementation. Bmpr2 expression remained relatively stable across groups, showing no significant changes with treatment. (**E**) Western blot of BMPR2 and loading control β -Actin. (**F**) Fiji ImageJ quantification analysis of BMPR2 normalized to β-Actin confirmed enhanced pathway activation in BMP9-treated grafts. Data are presented as mean ± SEM with individual replicates; P-values were determined by one-way ANOVA with multiple comparisons.

### Plasma from lung transplant recipients impairs pulmonary endothelial BMP9 signaling

Circulating levels of BMP9 decline in systemic inflammatory states such as sepsis and systemic inflammatory response syndrome (SIRS), yet BMP9’s functional relevance in the lung transplant setting remains undefined. Given the heterogeneous clinical manifestation of primary graft dysfunction (PGD) among transplant recipients, we hypothesized that impaired BMP9 signaling capacity prior to transplantation may stratify patients who go on to develop PGD. To evaluate this, pre-transplant plasma samples from lung transplant recipients were utilized and assessed for their capacity to induce BMP9 target gene expression, specifically ID1, in primary human pulmonary arterial endothelial cells (HAECs) and human lung microvascular endothelial cells (HMVEC-L) (Figure 6A). Clinical demographics of the patient cohort are summarized in Supplementary Table 1.

**Figure 6.**
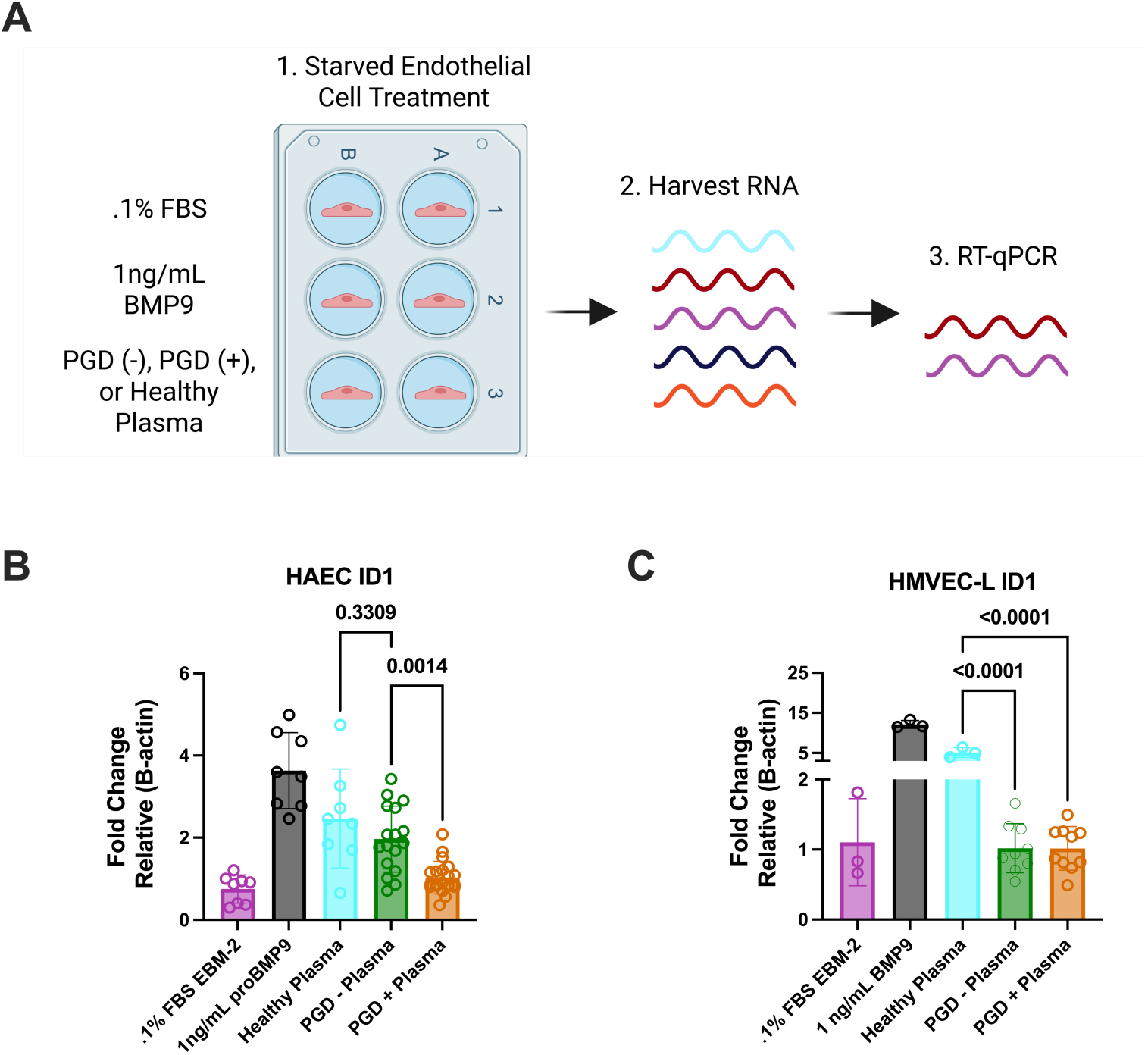
BMP9-induced transcriptional responses are impaired in plasma from lung transplant recipients. (**A**) Experimental schematic: human aortic endothelial cells (HAECs) and human microvascular endothelial cells (HMVEC-L) were serum-starved overnight (0.1% FBS) and treated with recombinant BMP9 (1 ng/mL), plasma from healthy controls, or plasma from lung transplant recipients with or without primary graft dysfunction (PGD). RNA was harvested and analyzed by RT-qPCR for BMP-responsive gene ID1. (**B**) In HAECs, BMP9 and healthy plasma robustly induced ID1 expression, whereas plasma from all LTx patients significantly blunted this response. (**C**) In HMVEC-L, both ID1 (**C**) and ID2 (**D**) were strongly induced by BMP9 and healthy plasma but suppressed in the presence of PGD+ plasma. Data are presented as fold change relative to β-actin, shown as mean ± SEM with individual replicates; P-values from one-way ANOVA with multiple comparisons are indicated.

Compared to healthy donor plasma, pre-transplant plasma from PGD+ patients significantly attenuated ID1 expression in HAECs, consistent with reduced circulating BMP9 bioactivity (Figure 6B). Notably, plasma from PGD– patients also impaired ID1 induction, albeit to a lesser degree, suggesting a broader suppression of BMP9 signaling in the lung transplant population as a whole. In HMVEC-Ls, which are highly responsive to BMP9 stimulation, plasma from both PGD– and PGD+ patients markedly blunted ID1 expression relative to healthy controls, further indicating a generalized impairment of BMP9 signaling in the pulmonary microvasculature (Figure 6C).

To assess whether the underlying disease etiology contributed to this signaling defect, we stratified patients by pre-transplant diagnosis. Interestingly, patients with interstitial lung diseases (ILDs) exhibited significantly reduced BMP9 signaling capacity compared to those transplanted for other indications, as shown in Supplemental Figure 4A & 4B. These findings are consistent with prior reports of BMP pathway dysregulation in chronic lung diseases and suggest that disease-specific alterations in BMP9 signaling may persist through transplantation. Collectively, these data reveal that circulating factors present in all lung transplant recipients regardless of PGD status are sufficient to impair endothelial BMP9 signaling prior to reperfusion.

Together with our murine data demonstrating rapid suppression of BMP9 signaling post-transplant, these results support the concept that exogenous BMP9 supplementation could serve as a broadly applicable therapeutic strategy to restore vascular homeostasis and mitigate early allograft injury.

## Discussion

Endothelial dysfunction is a hallmark of primary graft dysfunction (PGD), yet the molecular pathways responsible for maintaining endothelial stability during lung transplantation remain incompletely understood. In this study, we identify acute suppression of bone morphogenetic protein 9 (BMP9) signaling as a mechanistic feature of lung ischemia-reperfusion injury (IRI) and demonstrate that therapeutic BMP9 supplementation restores pulmonary endothelial signaling, attenuates immune cell infiltration, and improves graft function in a murine model of PGD. These findings establish BMP9 as a previously unrecognized regulator of vascular resilience in lung transplantation and support its translational potential as a therapeutic candidate.

BMP9 is a circulating ligand of the TGF-β superfamily that signals primarily through ALK-1 (ACVRL1) and BMPR2, a high-affinity receptor complex predominantly expressed on pulmonary endothelial cells (30). Canonical BMP9 signaling promotes vascular quiescence, endothelial barrier integrity, and anti-inflammatory responses through SMAD1/5/9 activation (31). Phospho-proteomics and of BMP9 signaling in pulmonary vascular endothelial cells revealed that BMP9 can regulate secondary signaling cascades including MKK4, P38, P27, and Notch (20, 32). These canonical and non-canonical functions of BMP9 signaling are particularly critical in the lung microvasculature, where injury-induced disruption of endothelial integrity leads to alveolar edema and impaired gas exchange (33). While previous studies have linked BMP9 pathway dysfunction to vascular disorders such as pulmonary arterial hypertension (PAH) and hereditary hemorrhagic telangiectasia (HHT), the role of BMP9 in acute sterile lung injury, particularly within the context of transplantation, has not been explored. Bleomycin induced lung sterile injury was shown to similarly negatively affect the BMP9/BMPR2/SMAD signaling axis (34).

Here, we show that transplantation-induced IRI rapidly downregulates BMP9 receptor components (*bmpr2, acvrl1*) and SMAD-dependent targets in murine lung allografts within six hours of reperfusion, consistent with a profound suppression of endothelial BMP9 responsiveness. These findings align with prior reports demonstrating that hypoxia, oxidative stress, and pro-inflammatory cytokines repress *bmpr2* and reduce BMP signaling in models of PAH and acute lung injury. Importantly, this disruption occurs in parallel with upregulation of inflammatory mediators such as Ccl2, Myd88, and Cxcl10, supporting the notion that BMP9 signaling functions to restrain early proinflammatory activation during IRI (35). Transcriptomic profiling revealed AQP1 as the most significantly downregulated gene at 24 hr following lung transplantation, yet the BMP9-BMPR2 signaling pathway was the focus of mechanistic studies because of its greater translational potential. Reduced AQP1 expression has been implicated in impaired alveolar fluid clearance and the development of pulmonary edema during lung transplant IRI, suggesting it plays a contributory role in barrier dysfunction and early graft injury. Unlike BMP signaling, AQP1 currently lacks selective and clinically viable pharmacologic modulators capable of enhancing its activity. In contrast, BMP9 signaling is a well-defined pathway central to pulmonary endothelial homeostasis, and recombinant BMP9 and related modulators are already in preclinical development. Focusing on BMP9 allowed us to interrogate a pathway with both a mechanistic role in microvascular integrity and high therapeutic tractability, which ultimately enabled us to test interventions that could translate more rapidly to clinical application.

Therapeutic administration of recombinant BMP9 at the time of reperfusion significantly improved arterial oxygenation, reduced histologic injury, and lowered circulating levels of soluble von Willebrand factor (svWF), a marker of endothelial activation (36). These protective effects are consistent with prior studies demonstrating that BMP9 limits endothelial permeability and leukocyte adhesion in inflammatory models (37, 38). Furthermore, BMP9-treated lungs exhibited significantly reduced neutrophil infiltration, suggesting that the vascular stabilization provided by BMP9 prevents early leukocyte extravasation into the graft. These data contrast that of previous studies showing increased endothelial cell immunogenicity in cytokine primed conditions highlighting the importance of temporal context in determining BMP9’s inflammatory effects (38, 39). In our models, administration of exogenous BMP9 following injury induced a therapeutic response and promoted a protective vascular phenotype, reducing susceptibility to PGD.

Mechanistically, BMP9-induced ID1 expression was enhanced at 6 hours post-transplant in the pulmonary vasculature without a corresponding increase in *bmpr2* transcription, suggesting that ligand availability, not receptor abundance is rate-limiting. These observations were further supported by in vitro findings in human pulmonary endothelial cells, where BMP9 stimulation increased BMPR2 protein levels despite unchanged mRNA, consistent with post-transcriptional receptor regulation (40). These are consistent with other reports that endothelial stimulation with BMP9 triggers BMPR2 protein expression (39, 40). Collectively, these findings indicate that early ligand supplementation is sufficient to restore signaling through pre-existing receptor pools, reinforcing the therapeutic rationale for BMP9 administration during the acute reperfusion phase, summarized in Figure 7.

**Figure 7.**
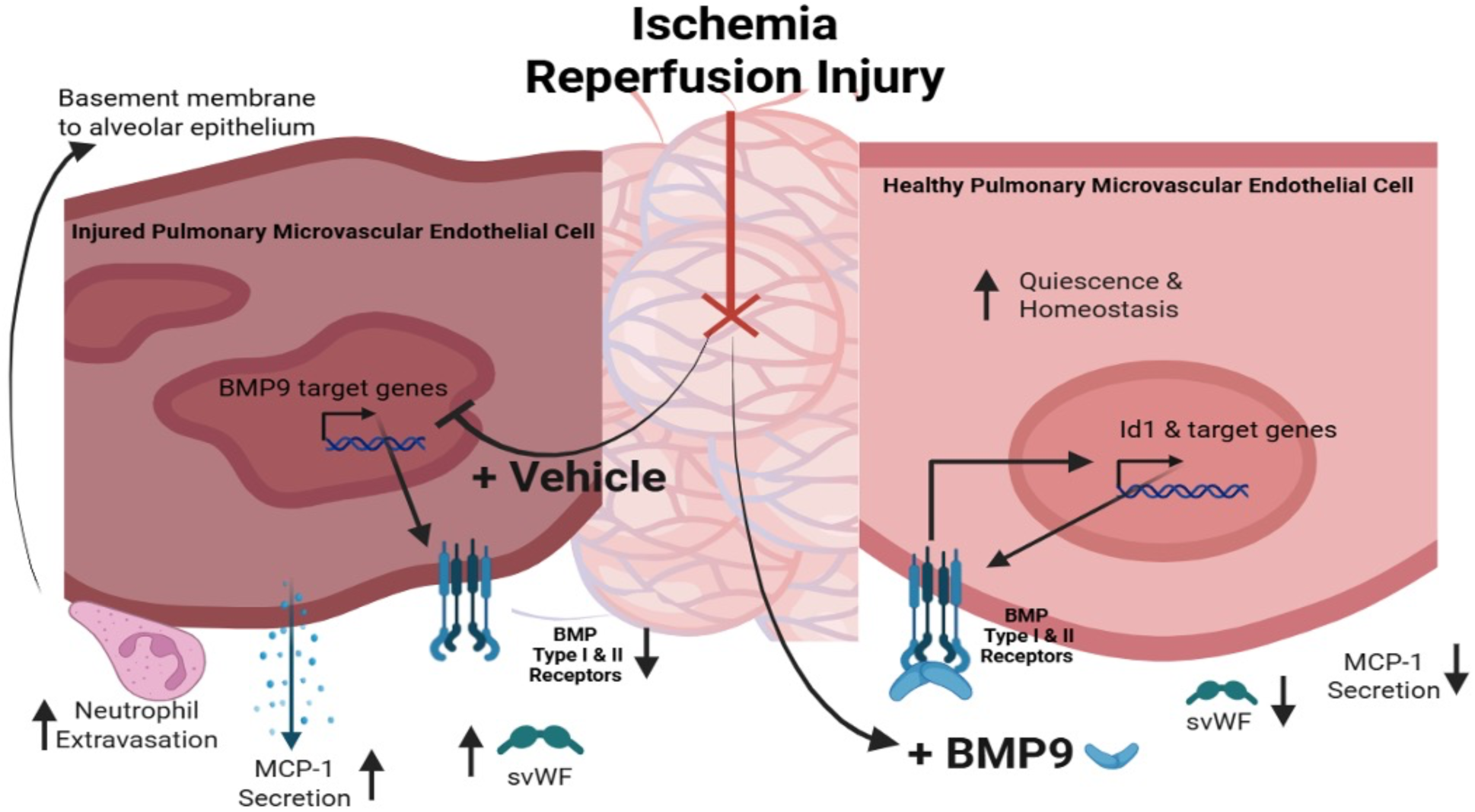
Graphical Abstract of how exogenous BMP9 therapy protected, and promoted a quiescent and an anti-inflammatory response, within the pulmonary microvasculature. BMP9 promoted robust induction of target gene expression, induced a quiescent phenotype, and dampened inflammation and injury.

To assess the functional relevance of these findings in human disease, we examined pre-transplant plasma from lung transplant recipients and found that plasma from both PGD+ and PGD– patients impaired ID1 induction using a primary human pulmonary endothelial cells reporter assay (41). While this suggests a global reduction in BMP9 bioactivity prior to transplantation, the effect was most pronounced in recipients with interstitial lung disease (ILD), aligning with prior reports of BMP pathway dysregulation in fibrotic lung disorders (42). Although circulating BMP9 protein levels were not directly measured in this cohort, these findings are consistent with prior studies demonstrating functional suppression of BMP9 signaling in systemic inflammatory conditions such as sepsis and cirrhosis, even when total ligand levels are unchanged (41). One possible explanation for this apparent signaling deficiency is the presence of circulating inhibitors or increased shedding of BMP9 co-receptors, such as soluble endoglin (sENG), which is known to bind BMP9’s growth factor domain in circulation (43). sENG was not measured in this study as it is known to act in cell-specific mechanisms and has dichotomous roles in integrating TGF/BMP superfamily signaling (44, 45).

Although our primary focus was on the pulmonary endothelium, BMP9 receptors are also expressed on immune cells, including alveolar macrophages, where ALK-1 activity has been shown to be dispensable (46). Whether BMP9 modulates alveolar macrophage phenotype, efferocytosis, or crosstalk with neutrophils during lung transplantation remains an open question. The observed reduction in neutrophil infiltration following BMP9 treatment may reflect direct vascular stabilization or an indirect effect on myeloid cell recruitment and activation. Additional studies using lineage-specific deletion or fate mapping will be necessary to disentangle these potential mechanisms.

Id1 expression was observed to be elevated at 6 hours but not at 24 hours post-transplant, which is consistent with the rapid kinetics of SMAD-mediated transcription in the endothelium. BMP9 naturally circulates in concentrations much higher than receptor affinity, as it constantly acts on the endothelium. Thus, these data indicate that the therapeutic efficacy of supraphysiological exogenous BMP9 may be temporally restricted (47). This observation underscores the importance of early intervention and suggests that BMP9 may be most effective when administered at the time of reperfusion. While our murine model does not incorporate immunosuppression, prior studies have shown that calcineurin inhibitors such as tacrolimus (FK506) can enhance BMP signaling by disrupting FKBP12-mediated inhibition of type I receptors (48, 49). Thus, future translational work should evaluate the interaction between BMP9 therapy and standard immunosuppressive regimens, particularly in relation to timing and dosing.

## Conclusions

This study establishes BMP9 signaling suppression as a novel mechanistic contributor to early endothelial injury and primary graft dysfunction in lung transplantation. Therapeutic BMP9 supplementation restores vascular BMP signaling, reduces neutrophil infiltration, and preserves lung function. Functional impairment of BMP9 signaling by human pre-transplant plasma, particularly in patients with fibrotic lung disease, supports the broader relevance of this axis in clinical transplantation. Together, these findings provide compelling rationale for the development of BMP9-based therapies aimed at restoring endothelial homeostasis and improving early graft outcomes in all lung transplant recipients.

## Methods

### Cell culture

Primary human lung microvascular endothelial cells (HMVEC-Ls) were purchased from Lonza Biosciences (CC-2527). HMVEC-Ls were grown to the manufacturer’s instructions (in CC-3202). Primary human arterial endothelial cells (HAECs) were purchased from Lonza Biosciences (CC-2535) and cultured to the manufacturer’s instructions (in CC-3162). The lot numbers and donor information for all primary cells can be found in Supplementary Table 2. HuLEC-5a were purchased from ATCC (CRL-3244) and grown in MCDB 131 (Gibco #10372019) supplemented with heat-inactivated fetal bovine serum, recombinant human EGF, hydrocortisone, L-Glutamine (Fisher #16140071, Corning #CB-40052, ThermoFisher #352450050, Gibco #25030081). All cells were cultured in penicillin/streptomycin (Gibco #15-140-122).

### BMP9 Activity Serum Starved Assay

HMVEC-Ls and HAECs were purchased from Lonza and maintained in 5% FBS per the manufacturer’s instructions. Prior to experimentation, cells were serum starved overnight, treated with 2% patient plasma, and harvested for mRNA for RT-qPCR. HMVEC-L mRNA was collected 5 hrs after patient plasma treatment, and HAEC mRNA was collected 1 hr after patient plasma treatment. These methods are adopted from previously published methods ^1^.

### RT-qPCR and NanoString

Qiagen RNeasy Mini Kit (Qiagen #74106) was used per the manufacturer’s instructions for mRNA extraction of mouse lung, HMVEC-L, and HuLEC-5a, respectively. All mRNA samples were treated with DNase I (Qiagen # 79256). For all in vitro and in vivo gene quantification studies, 1ug of mRNA was converted into cDNA using Applied Biosystems High-Capacity cDNA kit (ThermoFisher # 4368813). A QuantStudio 3 and Maxima SyBR Green qPCR Master Mix (ThermoFisher #K0252) was used for cDNA quantification. Mouse and human RT-qPCR primers are listed in Supplementary Table 3.

### NanoString nCounter Assay

Mouse left lung mRNA was harvested for NanoString nCounter Analysis. The mouse host response nCounter panel utilized 100ng of mRNA from each sample to quantify gene expression. nSolver 4.0. was used for gene normalization, advanced analysis, statistical workup, and data organization.

### In vitro cold storage ischemia reperfusion model

A modified cell culture model that simulates the cold storage, hyperoxia, and reperfusion process of LTx was used as previously described (50, 51) using HMVEC-L and HuLEC-5a cells, respectively. To test the impact of BMP9 treatment on endothelial injury BMP9 was added immediately post reperfusion. HuLEC-5a cell culture supernatant was harvested at 6 and 24 hrs post reperfusion for quantification of MCP-1 and IL-8 by ELISA (BD #555179, BD #555244).

### RNAscope

FFPE sections were cut, and the slides were dried at 60C for 1 hr. Slides were placed in xylene and 100% ethanol before air drying. Hydrogen peroxide was used to block slides before undergoing protease treatment. Advanced Cellular Diagnostic (ACD) Bmpr2 (#493061), Id1 (#312221), or control mRNA specific probes (UBC #310771, Negative Control #310043) were incubated on the slides which were then placed in the 40C ACD HybEZ II oven. Next, slides underwent 6 stages of amplification before being stained. Imaging for was conducted on a Keyence BZ-X810 with the BZ-X800 Viewer and BZ-X800 Analyzer applications. Five images from each sample were objectively scored on a scale from 0-3 by a blinded histologist for both Bmpr2 and Id1. The average score for each sample was plotted for each biological replicate.

### Immunohistochemistry

Tissue sections were cut and placed on slides before they were deparaffinized and rehydrated. Antigen retrieval was then performed with IHC TEK epitope retrieval solution (IHC World #IW11001L) and followed by a 1 hr block of goat serum (Sigma #G9023-5ML). Primary antibodies were incubated at 4C overnight. After a peroxide block, slides were washed in DAKO buffer (Agilent #S300685-2C) before 30min of secondary antibody exposure and DAB (Vector #SK-4100). Antibodies are listed in Supplementary Table 4.

### Immunofluorescence

Tissue sections were cut and placed on slides before they were deparaffinized and rehydrated. Slides were then added to coplin jars with IHC TEK epitope retrieval solution. Slides were then washed, blocked with Image-iT FX Signal Enhancer (ThermoFisher #I36933) for 30 min, washed, and blocked with goat serum for 20 min (Vector Laboratories #S101250). Primary antibodies were incubated at 4C overnight. On day 2, slides were washed, and secondary antibody was added for 45 min at room temperature. Afterwards, slides were washed and counterstained with DAPI (Vector Laboratories #H180010). Primary and secondary antibodies are listed in Supplementary Table 4.

### Western Blot

Cells were lysed with mammalian protein extraction reagent, M-PER (ThermoFisher #78501) containing 1X of HALT protease and phosphatase inhibitor (ThermoFisher #78441). Cellular debris was removed via centrifugation prior to protein quantification via Pierce BCA (ThermoFisher #23225). M-PER and 4x sample Buffer (ThermoFisher #NP0007) were used for sample preparation. Samples were loaded into Nupage Bis-Tris 4-12% gradient gels (ThermoFisher #WB14020) and ran wth NuPage MES-SDS running buffer (ThermoFisher #NP0002-02). Precision plus protein dual color standards (BioRad #1610374), Blotting Grade Blocker Non Fat Dry Milk (BioRad #1706404XTU), and TBS-T .1% were used in conjunction with the manufacturer’s protocol. Chemiluminescent detection utilized Western Lightning ECL (Revvity #NEL105001EA) and was performed on a ThermoFisher iBright Imager. For antibodies and dilutions, please see Supplementary Table 4.

### Lung transplantation

Male BALB/c and C57BL/6 mice were purchased from Jackson Laboratories. Eight-to twelve-week-old animals between 25 and 30g were used and housed in the specific pathogen free facility at University of Florida. All animal procedures were performed according to institutional animal care guidelines of the University of Florida. To evaluate the impact of BMP9 signaling on lung transplant outcomes, four experimental groups were designated for this study group: 1, Vehicle control; 2, BMP9 (1.5µg/kg); 3, anti-BMP9 antibody (5mg/kg); and 4, Isotype control antibody (5mg/kg). C57Bl/6 recipient mice were pretreated with either anti-BMP9 antibody or the isotype control 48 hr prior to LTx as well as immediately post LTx. Mice were euthanized 6 or 24 hours post-LTx for graft analysis and serum collection.

### Outcome Measures

Lung injury post transplantation was assessed using multiple domains, in line with the new ATS consensus report (52).

### Histological Analysis

To assess histological evidence of injury, lungs (native and Tx) were inflated en-bloc with formalin via the trachea using gravity inflation at 25cm H_2_O. Lung sections were stained with hematoxylin and eosin and scored for evidence of lung ischemic injury as previously described (53).

### Lung Function

At the end of the reperfusion period after LTx, mice were anesthetized, intubated, and placed on mechanical ventilation (10 μl/g tidal volume, respiratory rate 120/min, PEEP 3 cm H2O) with 100% oxygen for 5 minutes. Blood was collected from the left ventricle into heparinized syringes for arterial blood gas measurements. Arterial blood gases were measured using a VetScan i-STAT 1 handheld analyzer and CG4+ cartridges (Abaxis, Union City, CA).

### Lung Inflammation

To assess lung inflammation, innate immune cell influx was quantified by immunostaining with anti-MPO antibodies and quantified as previously described (54).

### Statistical Analysis

nSolver advanced analysis was used to generate volcano plots. Sartorius Simca 18.0. was used to perform PCA-X, hierarchal clustering, and Bi-Plot analyses. GraphPad Prism 10 was used to perform one-way ANOVA’s and student’s t test analyses, respectively.

## Data Availability

Supporting Data Values for all data presented in the manuscript and supplement can be found in Supplemental File 1. Raw Nanostring nCounter data can be found in GEO under accession GSE309366.

## Notes

https://www.ncbi.nlm.nih.gov/geo/query/acc.cgi?acc=GSE309366

### Competing Interest Statement

The authors have declared no competing interest.

### Funding

- National Heart Lung and Blood Institute: 1F31HL176117-01
- National Institute of Allergy and Infectious Diseases: 5U01AI70075-04

