## Supplemental Data File 1 for "Exogenous BMP9 therapy ameliorates primary graft dysfunction post lung transplantation"

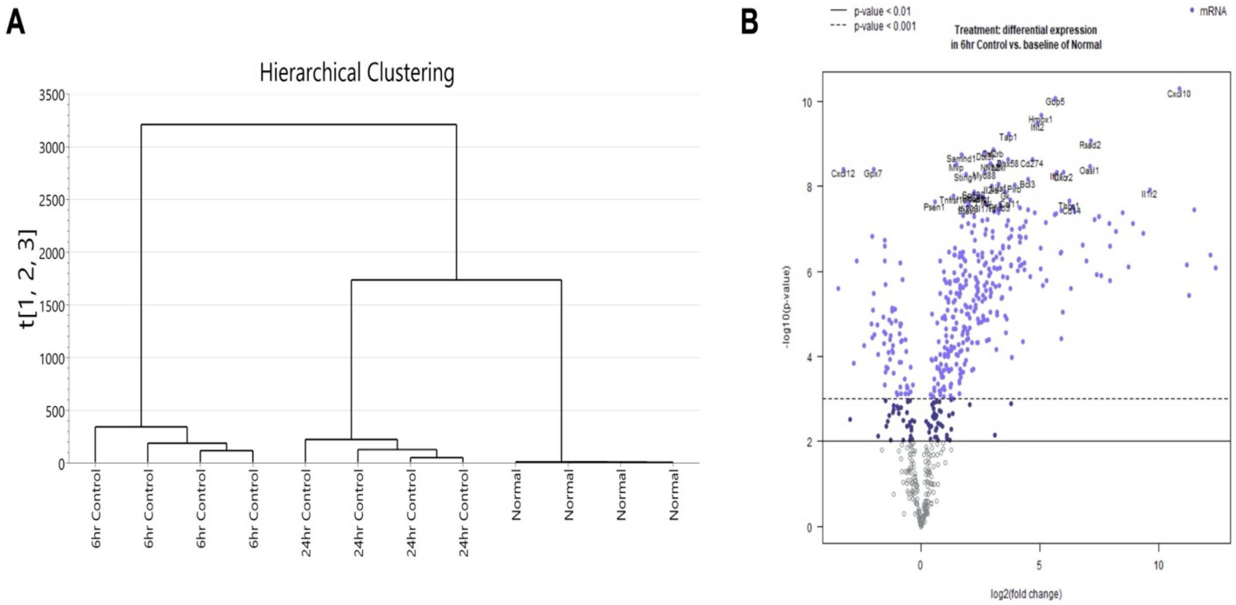


**Supplemental Figure 1. Hierarchical clustering reveals distinct transcriptomic separation of murine lung allografts post-transplant.** A. Nanostring nCounter normalized count data were analyzed by hierarchical clustering in Simca 18.0. Each gene was designated as a Y-variable and each biological replicate as an X-variable. Experimental groups (Normal, 6 hr post–cold storage ischemia-reperfusion injury (CS-IRI), and 24 hr post–CS-IRI) clustered distinctly, demonstrating reproducible segregation of replicates and highlighting unique transcriptional signatures at each time point following transplantation. B. nSolver generated volcano plot depicting differential gene expression between normal donor lungs and 6-hour post-transplant grafts. Genes were excluded from analysis if they were insufficiently expressed (average total count is under 100).

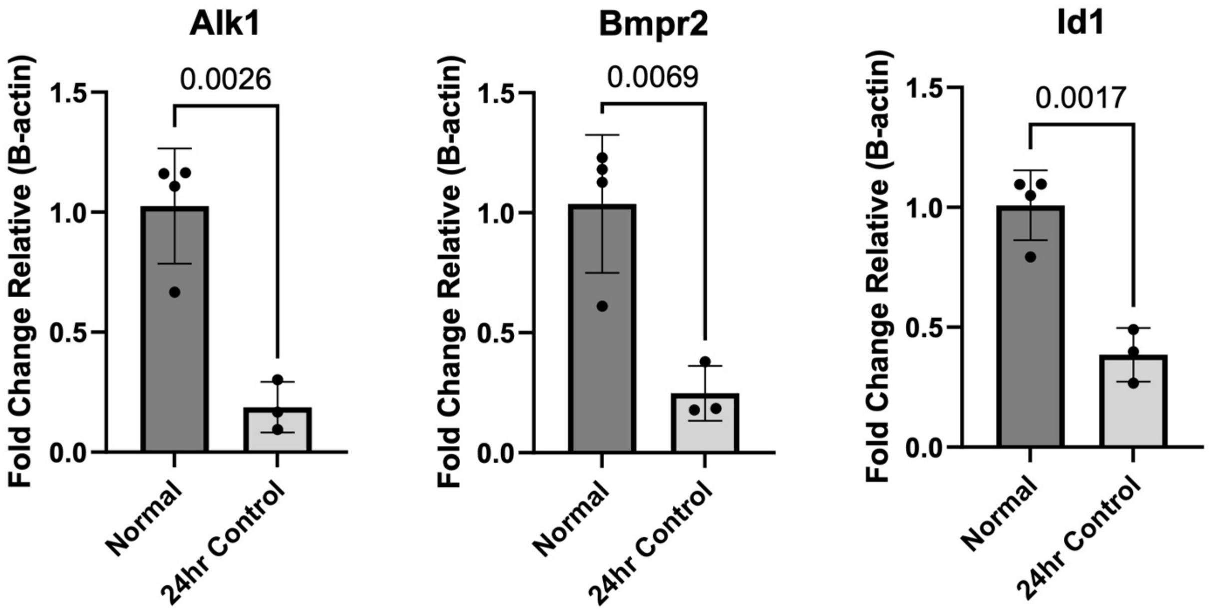


**Supplemental Figure 2.** RT-qPCR confirmation of Nanostring differential expression in murine left lung allografts post-transplantation. Gene expression levels of Acvrl1 (**A**), Bmpr2 (**B**), and Id1 (**C**) were measured and normalized to β-actin. Student's unpaired t-test was used; P-values indicated on each panel. Error bars represent mean ± SEM.

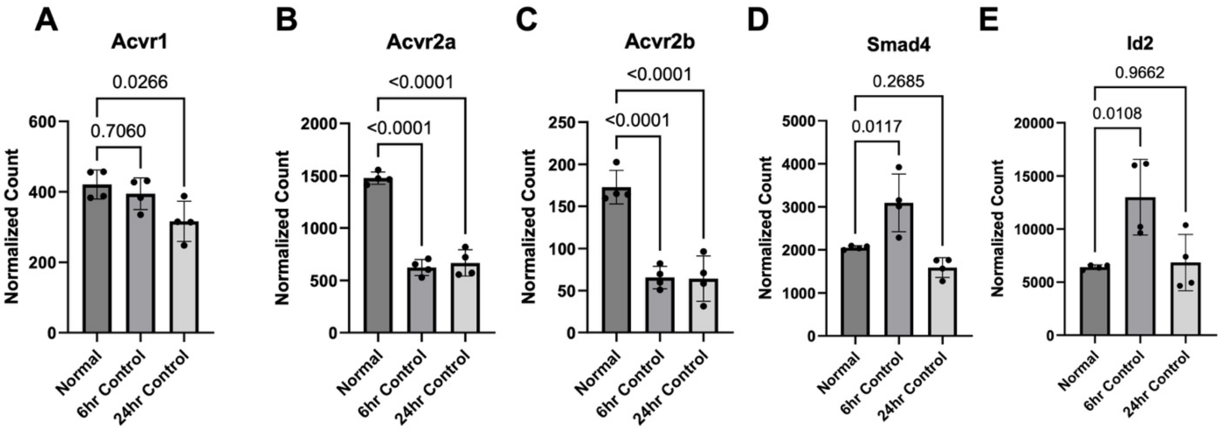


**Supplemental Figure 3.** Differentially regulated Nanostring nCounter BMP9 signaling genes. Bulk lung tissue expression of murine Acvr1, Acvr2a, Acvr2b, Smad4, and Id2. Student's unpaired t-test was used; P-values indicated on each panel. Error bars represent mean ± SEM.

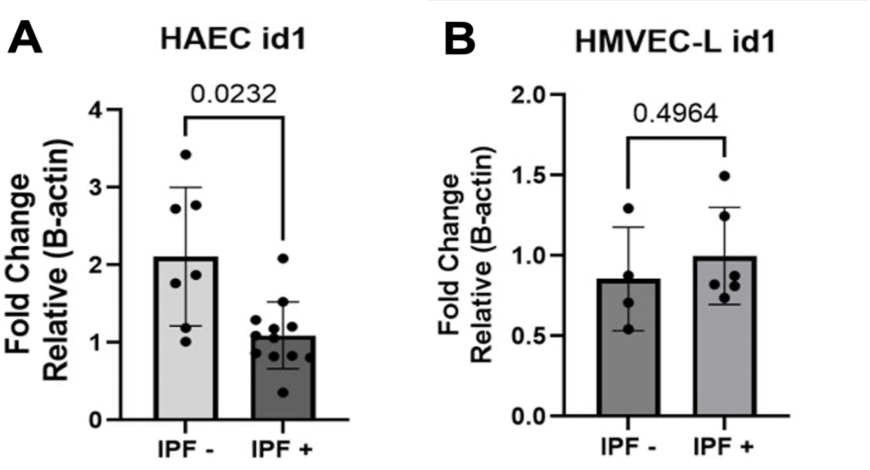


**Supplemental Figure 4.** Sub-analysis of pre-transplant diagnosis demonstrated impaired BMP9 signaling in IPF patients that developed PGD. Student's unpaired t-test was used; P-values indicated on each panel. Error bars represent mean ± SEM.

| | Non-PGD (n=9) | PGD (n=10) |
|---|---|---|
| **Recipient** | | |
| **Male sex, n (%)** | **7 (77.8)** | **4 (40.0)** |
| **Female sex, n (%)** | **2 (22.2)** | **6 (60.0)** |
| **Age in yr, median (IQR)** | **63 (61, 64)** | **56 (47, 62)** |
| **Diagnosis, n (%)** | | |
| **Idiopathic Pulmonary Fibrosis** | **4 (44.4)** | **6 (60)** |
| **COVID-19 Pulmonary Fibrosis** | **0 (0.0)** | **3 (30)** |
| **Chronic Obstructive Pulmonary Disease** | **2 (22.2)** | **1 (10.0)** |
| **Hypersensitivity Pneumonitis** | **1 (11.1)** | **0 (0.0)** |
| **Congenital Malformation** | **1 (11.1)** | **0 (0.0)** |
| **Bronchiectasis** | **1 (11.1)** | **0 (0.0)** |
| **Unknown** | **0 (0.0)** | **0 (0.0)** |

**Supplemental Table 1:** Lung Transplantation Patient Cohort Demographics. Patients without Primary Graft Dysfunction at any point post-LTx (PGD -, n=9) were compared to patients with primary graft dysfunction at all timepoints post-LTx (PGD+, n=10).

| Cell Type | Lot Number | Age (yr) | Gender | Race |
|---|---|---|---|---|
| **HMVEC-L** | 22TL052754 | 48 | Male | Caucasian |
| **HMVEC-L** | 22TL024424 | 43 | Female | Caucasian |
| **HMVEC-L** | 21TL316174 | 38 | Female | Caucasian |
| **HMVEC-L** | 22TL164370 | 50 | Female | Caucasian |
| **HMVEC-L** | 22TL238343 | 53 | Female | Caucasian |
| **HAEC** | 20TL231227 | 36 | Female | Other |
| **HAEC** | 20TL122171 | 56 | Male | Caucasian |

**Supplemental Table 2:** Donor Information for adult human microvascular lung endothelial cells (HMVEC-L) and adult human aortic endothelial cells (HAEC).

| Species | Target | Forward 5’ à 3’ | Reverse 5’ à 3’ |
|---|---|---|---|
| Human | ID1 | CTGCTCTACGACATGAACGGC | TGACGTGCTGGAGAATCTCCA |
| Human | BMPR 2 | CAAATCTGTGAGCCCAACAGT CAA | GAGGAAGAATAATCTGGATAAG GACCAAT |
| Human | ACVR L1 | CCATCGTGAATGGCTCGT | GGTCATTGGGCACCACATC |
| Human | SMAD 8/9 | ACACCACCCCTGCCTTATCA | CCTGGAATGTCTCCCCAACTC |
| Human | B-actin | CCCTGGACTTCGAGCAAGAG | CCAGGAAGGAAGGCTGGAAG |
| Mouse | Acvrl1 | CCAATGACCCCAGTTTTGAG | CTTTGGGCTTCTCTGGATTG |
| Mouse | Bmpr2 | AGCACAGAGGCCCAATTCTC | TCACCTATCTGTATACTGCTGCC |
| Mouse | Id1 | TGTTACTCACGCCTCAAGG | AACTGAAGGTCCCTGATGTAG |
| Mouse | B-actin | TACGTAGCCATCCAGGCTGT | GGAGAGCATAGCCCTCGTAG |

**Supplemental Table 3:** RT-qPCR Primers. Primers used to investigate gene expression in bulk murine lung tissue as well as human microvascular lung microvascular endothelial and aortic endothelial cells (HMVEC-L and HAEC, respectively).

| Application | Manufacturer | Catalog | Biologic | Dilution |
|---|---|---|---|---|
| **Murine Lung Transplant** | R&D Systems | MAB3209-500 | Anti-BMP9 antibody | 5mg/kg |
| **Murine Lung Transplant** | R&D Systems | MAB004 | IgG2B Isotype Control Antibody | 5mg/kg |
| **Murine Lung Transplant** | R&D Systems | 9624-BP-025 | Recombinant Latent Human BMP9 | 1.5 ug/kg |
| **In vitro IRI model** | R&D Systems | 9624-BP-025 | Recombinant Latent Human BMP9 | 1 ng/mL |
| **ELISA** | Abcam | ab208980 | Mouse vWF | |
| **Murine IF** | Abcam | ab9535 | MPO antibody | 1:300 |
| **Murine IF MPO** | ThermoFisher | A-32732 | Goat anti-rabbit 555 | 1:500 |
| **Murine IF P-SMAD1/5/9** | ThermoFisher | A-32732 | Goat anti-rabbit 555 | 1:200 |
| **BMPR2 Western Blot** | ThermoFisher | MA5-15827 | Anti-BMPR2 antibody | 1:500 |
| **B-actin Western Blot** | Cell Signaling Technologies | 12620 | Anti-B-actin-HRP | 1:1000 |
| **Detection – Western Blot** | Abcam | ab6728 | Anti-mouse-IgG-HRP | 1:5000 |

**Supplemental Table 4:** Biologics. Manufacturer and usage information for recombinant proteins, antibodies, and ELISA.

### References
